# Identification of small molecule enhancers of NK cell tumoricidal activity via a tumor microenvironment-mimicking co-culture assay

**DOI:** 10.1101/2024.09.04.611205

**Authors:** Aylin Binici, Elisabeth Hennes, Sandra Koska, Jens Niemann, Alisa Reich, Christiane Pfaff, Sonja Sievers, Astrid S. Kahnt, Dominique Thomas, Slava Ziegler, Carsten Watzl, Herbert Waldmann

## Abstract

The tumor microenvironment (TME) is a pro-cancerous niche harboring immunosuppressive factors that are secreted by cancer cells and the surrounding cancer-supportive tissue, such as kynurenine, prostaglandin E2 and transforming growth factor β (TGFβ). These factors dampen the activity of cytotoxic lymphocytes like natural killer (NK) cells, allowing evasion of immune cell-mediated killing. To identify small molecules that counteract the immunosuppressive effect of the TME and restore NK cell-mediated cytotoxicity, we developed a phenotypic co-culture assay of cancer cells and primary lymphocytes suitable for medium-throughput screening. We discovered small molecules that restore NK cell-mediated cytotoxicity through diverse mechanisms. The potent TGFβ type I receptor (TGFβR-1) inhibitor, RepSox, stood out as superior to other TGFβR-1 inhibitors due to its ability to abolish the effects of both inhibitory factors used in our setup. This mode of action goes beyond TGFβR-1 inhibition and is related to the simultaneous abrogation of cyclooxygenase 1 (COX1) activity.

## Introduction

Tumors escape immune surveillance through various mechanisms, including the activation of immunosuppressive signaling pathways and changes in metabolism.^1–3^ The tumor microenvironment (TME) is a heterogeneous assembly of various cell types such as cancer-associated fibroblasts, vascular endothelial cells, and immune cells like macrophages, dendritic cells, cytotoxic T lymphocytes, and natural killer (NK) cells.^4^ Cancer cells and the cancer-supportive tissue within the TME can gradually reprogram adjacent immune cells by releasing suppressive signaling factors such as adenosine, kynurenine (kyn), prostaglandin E2 (PGE_2_), and transforming growth factor β (TGFβ). This process leads to the adoption of an exhausted immunophenotype and a halt in anti-tumor responses, ultimately creating a pro-tumorigenic niche.^4–6^

In recent years, advanced knowledge of the TME architecture enabled the development of immunotherapies that enhance host immune response by overcoming suppressive mechanisms of the TME, thereby preventing immune exhaustion. This includes immune checkpoint inhibitors, like anti-PD-1 antibodies, and T cell-based therapies, such as genetically engineered CAR T cell treatments.^7,8^ Within the body, cytotoxic NK cells recognize and eradicate aberrant cells without prior sensitization such that NK cell-based therapies have been proposed for the treatment of cancer.^9–11^ Thus, the use of small-molecule modulators that inhibit immunosuppressive pathways or stimulate activating pathways may be a promising strategy for boosting the anti-cancer immune response of NK cells.^6,10^ However, previously reported assays employing co-cultures of tumor cells and NK cells have shown limited success in identifying potent small molecule enhancers for several reasons.^12–16^ This includes short incubation times with small molecules, which restricts hit identification to fast-acting compounds.^15^ Current methods for monitoring the target cell count are tied to endpoints and pose challenges such as the requirement to pre-stain cancer cells with fluorescent dyes such as calcein. This brings the risk of diluting fluorescence intensity due to target cell proliferation and false-positive hits from spontaneous releases.^16^ Release and detection of nano luciferase (NanoLuc) upon cancer cell lysis was employed for the detection of NK cell activity. However, this approach is limited to short assay times due to protein stability issues and small molecule-based luminescence interference.^12^ One of the main challenges for NK cell-based therapies targeting solid tumors is that NK cells must display optimal cytotoxic activity and viability to be effective in the suppressive TME. However, upon reaching the tumor site, the NK cell response is hindered due to suppressive factors in the TME, leading to exhaustion.^17,18^ Therefore, it is crucial for screening approaches to reflect the immunosuppressive nature of the TME, which has not yet been addressed in screening assays, along with optimal target cell monitoring over longer incubation periods.

To address the described limitations, we developed a novel TME-mimicking phenotypic co-culture assay for the identification of small molecule enhancers of NK cell cytotoxicity. The assay employs lung carcinoma A549 cells, which are susceptible to NK cell-mediated cytolysis, in co-culture with human donor-derived primary lymphocytes supplemented with the activating cytokine interleukin-15 (IL-15) as well as the immunosuppressive factors TGFβ and PGE_2_. This co-culture assay is amenable to medium-throughput screening and has identified several small molecules that enhance cancer cell cytolysis by NK cells operating by different mechanisms of action. The assay is broadly applicable to the discovery of novel modulators of NK cell cytotoxicity, restoring NK cell activity in the presence of immunosuppressive cytokines. The TGFβ type I receptor (TGFβR-1) inhibitor RepSox stood out as it efficiently enhanced NK cell cytotoxicity and eliminated the immunosuppressive effect of both inhibitory factors in our assay setup. This activity is attributed not only to the inhibition of TGFβR-1, and we demonstrate that RepSox is also a cyclooxygenase 1 (COX-1) inhibitor. This dual mode of action renders RepSox superior to all identified hit compounds and opens new avenues to the reactivation of NK cells in the TME.

## Results and Discussion

For assay development, the human lung adenocarcinoma cell line A549, which is susceptible to NK cell cytolysis, was stably transfected with an eGFP-tagged histone H2B type 1-J construct to yield A549^Green^ cells.^19^ This allowed monitoring the cytotoxic activity *via* the fluorescently labeled A549^Green^ nuclei in real time by means of kinetic live-cell imaging. IL-15 was selected to activate NK cells. Unlike frequently utilized IL-2, IL-15 stimulates NK cells without activating regulatory T cells (Treg), which can suppress the immune system and therefore is a disadvantage of IL-2-based adoptive immunotherapies.^20,21^ PGE_2_ and TGFβ were employed to mimic the immunosuppressive nature of the TME. TGFβ is one of the most powerful immunosuppressive cytokines, and is produced by a wide range of tumors to promote cancer immune evasion.^22^ TGFβ counteracts the IL-15-driven NK cell effector function by repressing mTOR signaling.^23^ PGE_2_ suppresses NK cell activity through prostaglandin E2 receptors EP2 and EP4 *via* the activation of adenylate cyclase, resulting in elevated intracellular cAMP levels.^24,25^ Both TGFβ and PGE_2_ were shown to decrease the expression of essential activating surface receptors such as NKG2D and NKp30 and simultaneously promote the expression of inhibitory surface receptors on NK cells.^26^ Since isolation of primary NK cells is demanding and provides only small amounts of suitable material, thereby limiting screening throughput, we resorted to primary lymphocytes to establish the screen. Primary lymphocytes were isolated from healthy donor-derived populations of peripheral mononuclear blood cells (PBMCs), which contain NK cells as well as T and B cells. Naïve cytotoxic T cells cannot initiate target cell cytolysis independently but require priming by antigen-presenting cells and cannot be activated solely by IL-15, while B cells do not exhibit any cytotoxic function.^27,28^ Therefore, when using lymphocytes under the described conditions, the reduction of A549^Green^ cell count should be exclusively mediated by NK cells. To detect NK cell-mediated tumoricidal activity, A549^Green^ cells and lymphocytes were directly co-cultured with the activating cytokine IL-15 alone or together with the immunosuppressive factors TGFβ and PGE_2_ (Figure 1A). The NK cell-mediated cancer cell cytolysis assay requires an incubation period of 120 to 144 h. This assay setup allows both, NK cell activation and compound treatment in a single well, saving time and enabling the identification of hits that have slower mechanisms of action.

**Figure 1:**
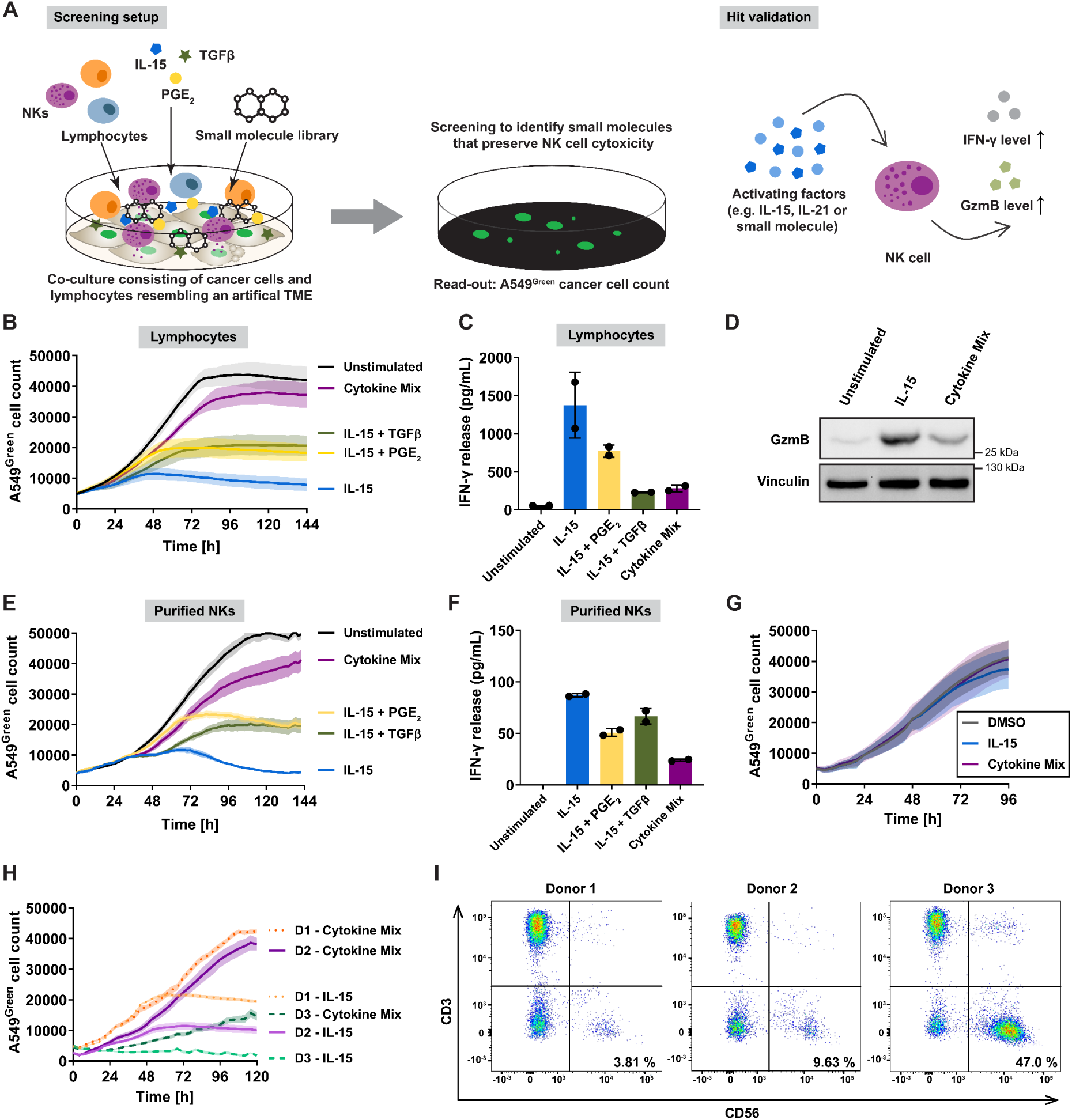
Development of an NK cell-mediated cancer cell cytolysis assay. A) Setup of the NK cell-mediated cancer cell cytolysis assay. A co-culture of A549^Green^ cells and lymphocytes or purified NK cells is treated with IL-15, TGFβ, and PGE2 and compounds. The cancer cell count is monitored for 120-144 h using the H2BjeGFP fluorescence. Cytokine mix: IL-15, TGFβ and PGE2. B) NK cell-mediated cancer cell cytolysis assay using primary lymphocytes. A549^Green^ cells were co-cultured with lymphocytes in the presence of different cytokines. The A549^Green^ cell count was monitored by means of live-cell imaging. Representative graph of n=3 (mean values ± SD, N=3 and E:T 60:1). E:T: effector-to-target ratio. The error bands represent SD. C) IFN-γ secretion of lymphocytes treated with different combinations of IL-15, TGFβ, and PGE2. IFN-γ levels were determined in supernatants of the NK cell-mediated cancer cell cytolysis assay using ELISA. Representative graph of n=3 (data are mean values ± SD, N=2). D) Influence of the assay conditions on the granzyme B (GzmB) levels. Lymphocytes were stimulated with IL-15 only or in presence of TGFβ and PGE2 in presence of DMSO for 48 h prior to the quantification of protein levels *via* immunoblotting. Representative blot of n=3 is shown (see also Figure S7). E) NK cell-mediated cancer cell cytolysis using purified NK cells. Representative graph of n=3 (mean values ± SD, N=3 and E:T 10:1). F) Secretion of IFN-γ by NK cells treated with IL-15 only or combined with TGFβ and PGE2. IFN-γ was detected in supernatants from the NK cell-mediated cancer cell cytolysis assay. Representative graph for n=3 (mean values ± SD, N=2). G) Influence of cytokines on A549^Green^ viability. A549^Green^ cells were treated with IL-15 alone or the cytokine mix. The cell count was monitored using the fluorescence of H2BjeGFP *via* live-cell imaging (mean values ± SD, n=3). H) Exemplary comparison of the assay windows of different donor lymphocytes. Cells were treated with IL-15 or the cytokine mix in the NK cell-mediated cancer cell cytolysis assay. Donor 1 (D1) E:T 40:1, Donor 2 (D2) E:T 40:1 and Donor 3 (D3) E:T 20:1. Data are mean values ± SD. I) Flow cytometry analysis of CD3 and C56 on lymphocyte donors used in H. The NK cell proportion can be found in the lower right quadrant (see Figure S1 for the gating strategy).

Initial investigation of the individual suppressive factors in the co-culture assay (Figure 1B) revealed that IL-15 initiated target cell lysis after approximately 48 h. TGFβ or PGE_2_ partially reduced NK cell cytotoxic activity induced by IL-15. The combination of IL-15, TGFβ, and PGE_2_ (referred to as cytokine mix) decreased the cytotoxic activity of NK cells to a level comparable to the unstimulated condition. This is in line with the additive inhibitory effect of TGFβ in combination with PGE_2_ on NK cell cytotoxicity.^29–31^ We validated the inhibitory effect of the suppressive factors on NK cell activity by monitoring IFN-γ production and the levels of apoptosis-inducing protein granzyme B (Gzm B).^32,33^ Exposure of lymphocytes to IL-15 elevated the levels of IFN-γ (Figure 1C), and IFN-γ secretion was reduced by the addition of either TGFβ or PGE_2_ individually or with the cytokine mix. Moreover, the GzmB level was increased upon IL-15 treatment and reduced by the cytokine mix (Figure 1D). These findings are consistent with the results of the cytolysis assay. Analysis of isolated NK cells displayed comparable trends, thereby validating the assay setup (Figures 1E and 1F). Neither IL-15 nor the cytokine mix impaired A549^Green^ cell viability, validating that the reduction of the A549^Green^ cell count is mediated by NK cell cytotoxicity (Figure 1G).

The fraction of NK cells in total PBMC populations can vary between 5 and 20% among different donors, calling for pre-evaluation of each donor material.^34^ Therefore, the optimal effector-to-target (E:T) ratio for each donor was determined by assessing the extent of A549^Green^ cell lysis in the presence of IL-15 compared to the cytokine mix to assure a viable assay window (Figure 1H). Moreover, we determined the proportion of CD3^-^ and CD56^+^ cells within donor lymphocytes, which represent the NK cell population, as CD3 is expressed on T cells, while CD56 is a marker for NK cells.^35^ Thereby, the NK cell proportion was related to the cytolysis efficiency of the respective donor (Figure 1l). Donor 1-derived lymphocytes contained 3.8% NK cells and displayed a lower cytotoxic activity using the same E:T of 40:1 compared to donor 2 with 9.6% NK cells. The sample from donor 3 contained 47.0% NK cells which is consistent with the high cytotoxic activity observed. Lymphocytes with unusually high NK cell proportions, such as donor 3, were excluded from further assays. Altogether, this finalized setup was utilized for the identification of small molecules that can either prevent the effect of the immunosuppressive factors or stimulate NK cell activating pathways.

A total of 29,502 compounds were screened in 1536 format at 11 µM (Z’-value: 0.3-0.4; signal-to-background ratio S/B: 2-14). Of those, 19,396 were commercial compounds selected from various sources including ChemDiv, the Edelris library (Keymical Collections™), the LOPAC® library, and the Prestwick Chemical Library® of FDA-approved and EMA-approved drugs. 10,106 compounds are part of our unique in-house library containing natural product (NP)-inspired compounds and pseudo NPs.^36,37^ As compounds may directly decrease the A549^Green^ cell count through toxicity, the influence of hit compounds on A549^Green^ cell growth was assessed in the absence of lymphocytes and additional factors. Hits that either decreased cell count by more than 25% or whose viability IC_50_ and cytolysis EC_50_ did not differ by a factor greater than four were excluded from further analysis. Dose-response analysis resulted in a final number of 11 hit compounds that increased NK cell-mediated cancer cell killing (Table 1).

**Table 1:** List of hit compounds that enhance NK cell cytotoxicity together with known targets. Automated cytolysis assay data are mean values ± SD, n=3. CHIR-99021 was tested at 10 µM while BIO was assayed at 11 µM. The influence of hit compounds on cell viability was evaluated using Hoechst-33342 staining to determine the cell count (*) or CellTiterGlo™ (CTG; **^#^**) assay after 144 h. Data are mean values ±S.D., N=3. Compounds highlighted in gray were excluded from further analyses due to cytotoxicity.

| Compound | Automated NK cell-mediated cell cytotoxicity assay<br>EC<sub>50</sub> [μM] or % activity | A549<sup>Green</sup> viability IC<sub>50</sub> via cell count* / CTG# [μM] | Mode of action |
| --- | --- | --- | --- |
| CHIR-99021 | 29% | > 17* | GSK3β inhibitors |
| BIO | 35% | n.d. |  |
| SAR405 | 0.74 ± 0.30 | 13.17 ± 1.83<sup>#</sup> | VPS34 inhibitor |
| VPS34-IN1 | 0.72 ± 0.22 | 6.77 ± 0.18* | |
| PIK-III | 1.30 ± 0.27 | > 6* | |
| Compound 19<br>PIK-III analogue | 1.60 ± 0.37 | > 6* | |
| MZ-1 | 0.13 ± 0.04 | 1.33* | PROTAC of BRD4, BRD2 and BRD3 |
| (+)-JQ1 | 0.14 ± 0.03 | 0.06* | Inhibitor of BRD4 (1/2) |
| BAY-299 | 0.30 ± 0.02 | 1.47* | Dual inhibitor of BRD1 and TAF1 |
| OTX015 | 0.34 ± 0.08 | 0.45* | Inhibitor of BRD2, BRD3, and BRD4 |
| Molibresib besylate | 0.37 ± 0.13 | 0.17* | BET bromodomain inhibitor |
| I-BET151 | 1.30 ± 0.13 | 3.10* | Pan BET family inhibitor |
| RepSox | 0.18 ± 0.03 | > 1.7* | TGF-β type I receptor inhibitors |
| SB-525334 | 0.33 ± 0.02 | 11.51 ± 2.16<sup>#</sup> | |
| SB-505124 | 0.47 ± 0.13 | >6<sup>#</sup> | |
| TP-008 | 0.98 ± 0.24 | >3<sup>#</sup> | |
| GW788388 | 1.19 ± 0.39 | >30<sup>#</sup> | |
| Galunisertib | 2.07 ± 0.46 | > 17* | |
| LY-2109761 | >3.4* | > 6* |  |

The screen identified the glycogen synthase kinase 3β (GSK3β) inhibitors CHIR99021 and 6-bromo-indirubin-3’-oxime (BIO) (Figure 2A) as hits. GSK3β inhibitors have been linked to NK cell biology as they enhance NK cell maturation, cytokine production, and tumoricidal activity.^38–40^ CHIR99021, and BIO significantly elevated the cytolysis rate by 75% and 53%, respectively (Figure 2B) in the presence of the suppressive cytokine mix. Compound-mediated NK cell activation was validated *via* the increased Gzm B levels for both inhibitors (Figure 2C and 2D). It has previously been demonstrated that both GSK3β inhibitors elevate IFN-γ production in NK cells.^38,41^ These results validate our assay setup as it identified small molecules previously reported to enhance NK cell cytotoxicity. G) Chemical structure of MZ-1. H and K) NK cell-mediated cancer cell cytolysis assay in the presence of MZ-1 using lymphocytes (H) and purified NK cells (K), respectively (mean values ± SD, n=3) ; *, P < 0.05; **, P < 0.01; ***, P < 0.001; ****, P < 0.0001 (one-way ANOVA, comparison to cytolysis rate of cytokine mix-treated control)). I and L) Influence of MZ-1 on the IFN-γ secretion of lymphocytes (I) and purified NK cells (L). Representative data of n=3 (mean values ± SD, N=2). J) Influence of MZ1 on the GzmB protein levels of lymphocytes after 48 h. Representative immunoblot (n=4, see also Figure S7).

**Figure 2:**
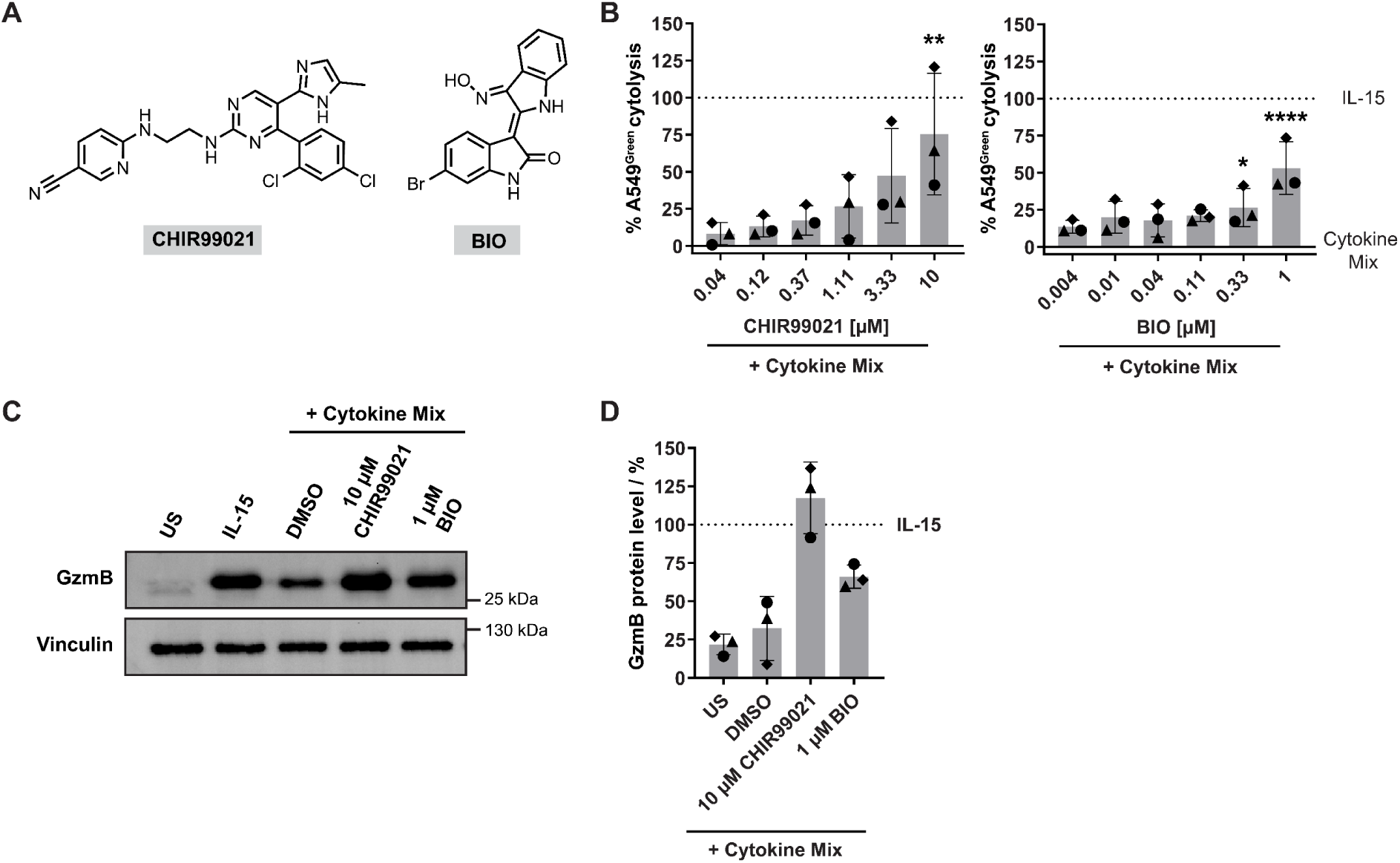
The GSK3β inhibitors CHIR99021 and BIO enhance NK cell cytotoxicity. A) Chemical structures of the GSK3β inhibitors CHIR99021 and BIO. B) NK cell-mediated cancer cell cytolysis in presence of CHIR99021 or BIO using lymphocytes (mean values ± SD, N=3, n=3) *, P < 0.05; **, P < 0.01; ***, P < 0.001; ****, P < 0.0001 (one-way ANOVA, comparison to cytolysis rate of cytokine mix-treated control)). C and D) Influence of CHIR99021 and BIO on the Gzm B levels of lymphocytes after 48 h of treatment. C) Representative immunoblot of n=3 (see also Figure S7). D) Quantified GzmB protein levels (mean values ± SD, n=3). US: unstimulated cells.

SAR405 and MZ-1 were identified as hits for which no link to enhanced NK cell tumoricidal activity is reported yet (Figure 3A and 3G). The phosphatidylinositol 3-kinase (VPS34) inhibitor SAR405 increased the cytolysis rate of lymphocytes and purified NK cells (Figure 3B and 3E).^42^ The compound elevated IFN-γ secretion (Figure 3C and 3F) and Gzm B levels in lymphocytes (Figure 3D). The lipid kinase VPS34 is essential for autophagy initiation and progression, and there is a link between autophagy and the degradation of Gzm B.^43–46^ Inhibiting VPS34 using SAR405 may suppress autophagy and impede Gzm B degradation, and this, in turn, may be the cause for the observed increase of NK cell-mediated cancer cell cytolysis. Additional VPS34 inhibitors, such as VPS-IN1, displayed an unsuitable window between A549^Green^ viability IC_50_ and NK cell cytolysis EC_50_ and were deemed unfit for deeper characterization (Table 1). The proteolysis targeting chimera (PROTAC) MZ-1 induces proteasomal degradation of bromodomain-containing proteins (BRD) 2,3 and 4.^47^ MZ-1 significantly enhanced the cytotoxic activity in the lymphocyte as well as the isolated NK cell setup in a concentration-dependent manner (Figure 3G and 3H). It dose-dependently upregulated IFN-γ secretion in lymphocytes as well as purified NK cells (Figure 3I and 3L). However, MZ-1 did not affect Gzm B levels in lymphocytes (Figure 3J). In MZ-1, the BRD2/3/4 inhibitor (+)-JQ1 and a VHL ligand derivative are linked. (+)-JQ1 displayed high toxicity in A549^Green^ (Table 1) while at non-toxic concentrations, the compound was inactive in the cytolysis assay (Figure S2). The derivative of the VHL ligand did not enhance NK cell cytotoxicity. Therefore, neither (+)-JQ1 nor the VHL ligand alone is responsible for the activity of MZ-1 in the co-culture assay (Figure S2). Different cellular outcomes occur after the inhibition of BRD4, depletion using RNA interference, or degradation using PROTACs.^48^ BRD4 inhibition is cytostatic, while BRD4 degradation *via* PROTACs is cytotoxic for cells. This is because the PROTAC-induced degradation leads to the generation of neo-tetrapeptides that deactivate Inhibitors of Apoptosis (IAP) proteins, which in turn increase caspase activation and thereby promote cell death through apoptosis. Active IAP signaling protects target cells from NK cell-mediated cytolysis, and thus, antagonizing IAP signaling may enhance susceptibility towards NK cell-mediated cytolysis.^49,50^ Other bromodomain inhibitors tested in the screening, such as BAY-299 and OTX015, showed as well an insufficient window between viability IC_50_ of A549^Green^ cells and NK cell cytolysis EC_50_ (Table 1).

**Figure 3:**
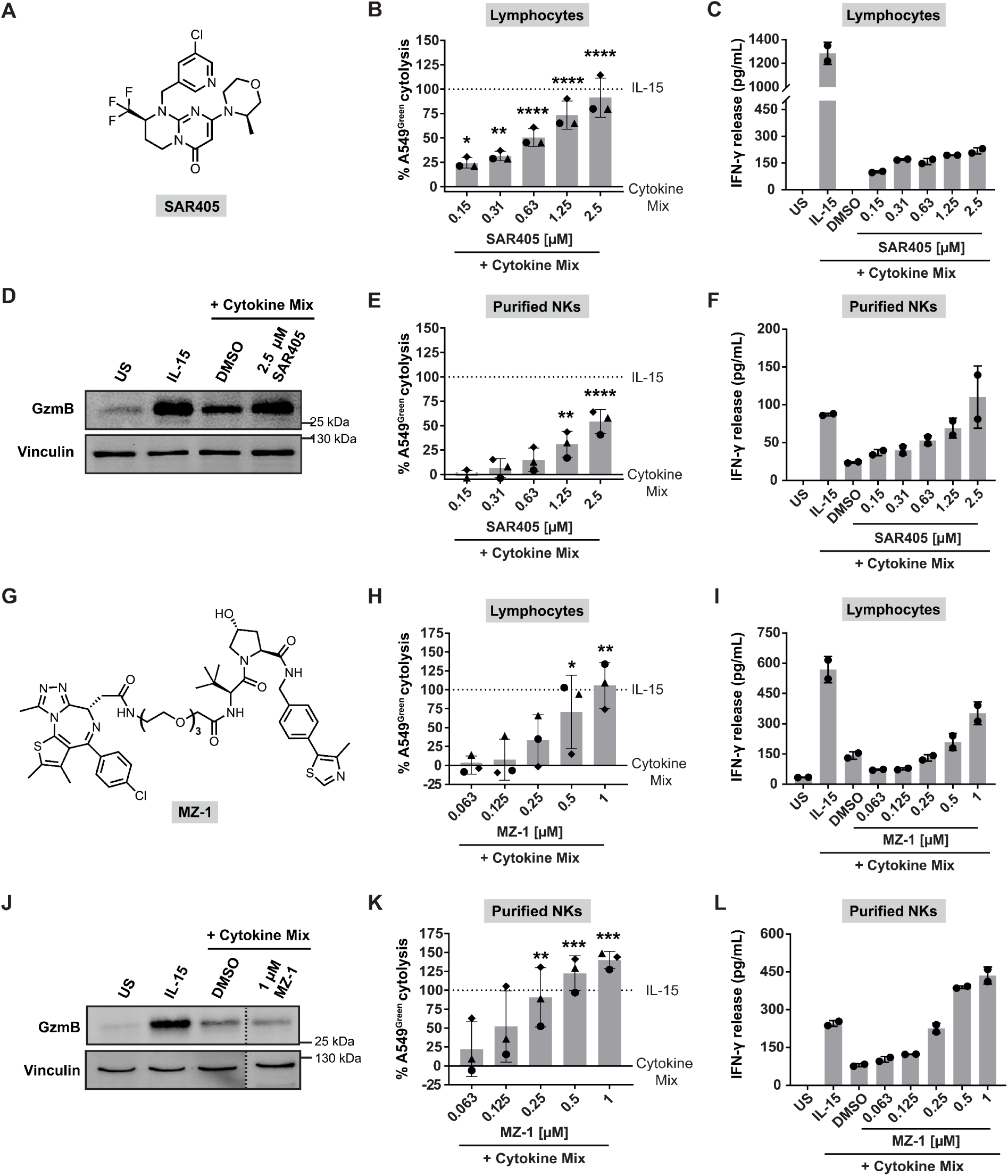
Validation of hit compounds SAR405 and MZ1. A) Chemical structure of SAR405. NK cell-mediated cancer cell cytolysis assay in presence of SAR405 using lymphocytes (B) and purified NK cells (E). Data are mean values ± S.D, n=3; *, P < 0.05; **, P < 0.01; ***, P < 0.001; ****, P < 0.0001 (one-way ANOVA, one-way ANOVA, comparison to cytolysis rate of cytokine mix-treated control)). C) and F) Influence of SAR405 on the IFN-γ secretion of lymphocytes and NK cells, respectively. Data are mean values ± SD (N=2) and representative of n=3. D) Influence of SAR405 on the GzmB protein levels after 48 h of treatment of lymphocytes. Representative immunoblot of n=3 (see also Figure S7).

Moreover, the screen identified the TGFβR-1 inhibitors RepSox, SB-525334, SB-505124, GW788388, LY2109761, and galunisertib (Figure 4) as hits, which was an expected finding since TGFβ was employed as inhibitory factor in the assay.^51–56^ Indeed, galunisertib can restore NK cell cytotoxicity and Gzm B levels in presence of TGFβ.^57,58^ For side-to-side comparison of the obtained NK cell cytolysis screening assay data, the identified TGFβR-1 inhibitors were subjected to a TGFβ-responsive Smad binding element (*SBE4*) dependent reporter gene assay (Figure 4B).^59^ We observed a positive correlation between the cytotoxic activity of NK cells and the inhibition of *SBE4*-dependent reporter gene expression. RepSox displayed the highest inhibition potency with an IC_50_ value of 8.5 nM and was the most effective enhancer of NK cell tumoricidal activity with an EC_50_ value of 180 nM. LY2109761 was the least effective in decreasing the *SBE4* reporter expression with an IC_50_ value of 861 nM. In line with these data, the cytotoxic activity of NK cells in the presence of LY2109761 was only weakly enhanced with an EC_50_ >3.4 µM. Analysis of NK cell-mediated cancer cell cytolysis, IFN-γ secretion, and Gzm B level in a manual setup revealed that the NK cytotoxic activity moderately parallels Gzm B levels (Figure 4C). However, there was no observable trend for IFN-γ secretion. Furthermore, 250 nM of RepSox was sufficient to enhance NK cell-mediated cytolysis to the level of IL-15-treated cells (Figure S4A). Notably, a potentiation of NK cell cytotoxicity beyond the level of IL-15 stimulation was detected for 1 µM RepSox with a cytolysis rate of 118%. IFN-γ secretion and Gzm B levels in RepSox-treated lymphocytes increased to 704% and 349%, respectively. Treatment of either lymphocytes or purified NK cells with the cytokine mix combined with a serial dilution of RepSox resulted in dose-dependent enhancement of the cytotoxic activity of NK cells (Figure 1D), revealing a similar EC_50_ value of 120 nM for lymphocytes and 104 nM for purified NK cells. RepSox is slightly more active in NK cell-mediated cytolysis and *SBE4* reporter gene assay than SB-525334. However, an increase in cytolysis rate, IFN-γ and GzmB levels exceeding the amount in IL-15-treated samples were not observed for treatment with SB-525334 or for the remaining TGFβR-1 inhibitors. These results indicate that all tested TGFβR-1 inhibitors besides RepSox most likely suppress only TGFβ signaling in the NK cell cytolysis assay. RepSox, however, counteracts the inhibitory effect of both TGFβ and PGE_2_ on cytolysis since the extent of cytolysis in the presence of the cytokine mix and 1 µM RepSox paralleled or even exceeded the cytolysis rate after IL-15 stimulation. Previous studies comparing the biological activity of TGFβR-1 inhibitors repeatedly demonstrated that RepSox activity goes beyond TGFβR-1 inhibition.^60,61^ Of note, all identified TGFβR-1 inhibitors have a very similar scaffold, and several compounds with this chemotype have common off-targets such as vascular endothelial growth factor receptors (VEGFR), receptor-interacting serine/threonine-protein kinase 2 (RIPK2), and mitogen-activated protein kinase 14 (p38α).^62–65^ To rule out any influence by off-targets of this chemotype, we investigated the highly selective TGFβR-1 and Activin-like kinase 4 (ALK4) inhibitor TP-008 (Figure 5A), which has a different scaffold.^66^ Hanke *et al*. profiled TP-008 in a panel of 469 kinases which revealed no additional kinase targets.^67^

**Figure 4:**
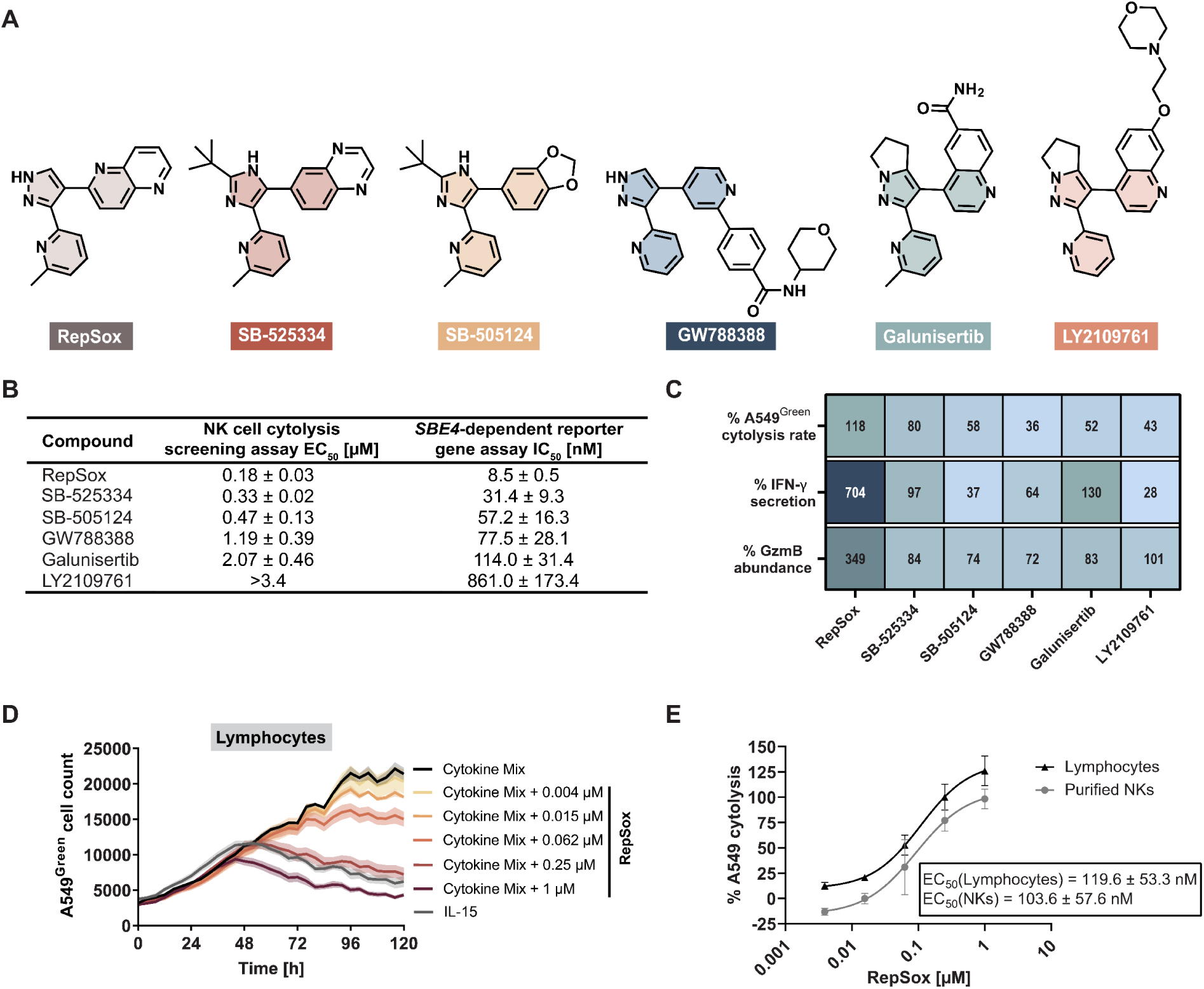
TGFβ receptor type I receptor (TGFβR-1) inhibitors enhance NK cell cytotoxicity. A) Chemical structures of RepSox, SB-525334, SB-505124, GW788388, galunisertib and LY2109761. B) Comparison between the cytotoxic activity in the automated NK cell-mediated cytolysis screening assay and the inhibitory activity in the *SBE4*-dependent reporter gene assay. Data are mean values ± SD (n=4 for RepSox, SB-525334 as well as galunisertib and n=3 for SB-505124, GW788388 and LY2109761, see also Figure S3). C) Heatmap representation of the activity of TGFβR-1 inhibitors RepSox (1 µM), SB-525334 (2 µM), SB-505124 (2 µM), GW788388 (20 µM), galunisertib (20 µM in the NK cell-mediated cytolysis assay and IFN-γ level and, 10 µM for GzmB protein level assessment), LY2109761 (10 µM) in the NK cell-mediated cancer cell cytolysis assay, the IFN-γ ELISA and Gzm B immunoblots using lymphocytes treated with the cytokine mix and the compounds (see also Figures S4-S5 and S7). The NK cell-mediated cytolysis assay data were normalized to IL-15 as 100% and cytokine mix as 0%. The IFN-γ, as well as Gzm B level data, were related to the levels in IL-15 treated cells (set to 100%). Mean values of n=3 for the cytolysis assay, IFN-γ, and Gzm B levels for all compounds except for SB-505124, for which Gzm B level data are mean values for n=4. D) NK cell-mediated cancer cell cytolysis assay in presence of RepSox using lymphocytes. Representative graph of n=3 (mean data ± SD, N=3, E:T = 40:1). E) Determination of the cytolysis rate of RepSox-treated lymphocytes and NK cells in the NK cell-mediated cancer cell cytolysis assay (mean values ± SD, n=3).

**Figure 5:**
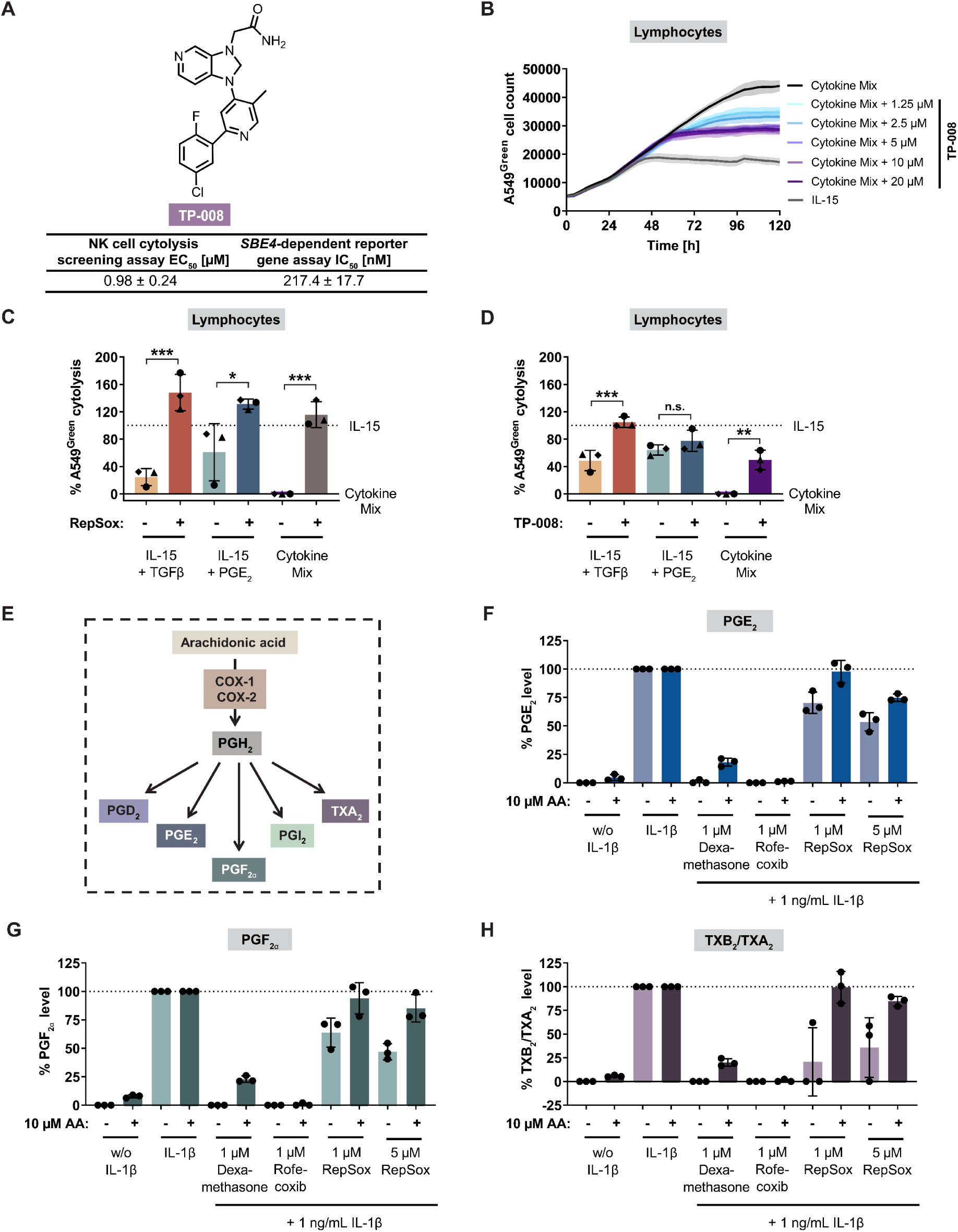
RepSox attenuates the immunosuppressive effect of TGFβ and PGE2. A) Chemical structure of the TGFβR-1 inhibitor TP-008. B) NK-cell mediated cancer cell cytolysis in the presence of TP-008 using lymphocytes with an E:T of 60:1. Representative graph (mean ± SD, n=3). C) and D) Influence of 1 µM RepSox and 20 µM TP-008 on the NK cell cytotoxic activity using lymphocytes. Data are mean values ± SD (n=3); P < 0.05; **, P < 0.01; ***, P < 0.001; ****, P < 0.0001, n.s.: not significant (two-way ANOVA). E) Cyclooxygenase (COX) 1 and 2 metabolize arachidonic acid (AA) into PGH2, which Is then further converted to PGD2, PGE2, PGF2α, PGI2, and TXA2 by several downstream synthases. F-H) Investigation of the influence of RepSox on the prostanoid and thromboxane production in A549 cells. Metabolite levels in presence or absence of AA were normalized to their respective IL-1β control as 100% (mean ± SD). The glucocorticoid Dexamethasone, inhibiting IL-1β-induced COX-2 expression, and reversible COX-2-selective inhibitor Rofecoxib were used as controls.

TP-008 enhanced the cytotoxic activity of NK cells (Figure 5B). However, even at 20 µM, it did not boost the cytotoxic activity to the extent of the IL-15 treatment. Hence, TP-008 cannot completely revert the influence of TGFβ and PGE_2_, indicating that it antagonizes TGFβ signaling only. In contrast to RepSox, TP-008 abolished only the effect of TGFβ on cytolysis, whereas the suppressive effect of PGE_2_ remained unaltered (Figure 5C-D). These results suggest that RepSox may modulate PGE_2_ signaling as well (Figure 5C and S4). Cyclooxygenases generate prostaglandins (PG), including PGE_2_ and thromboxanes (TX) from arachidonic acid (AA). Both COX isoforms contribute to carcinogenesis, and elevated PGE_2_ levels are often observed in cancer. They reduce anti-tumor immunity by decreasing the cytotoxic function of NK cells.^24,68^ Therefore, we analyzed the impact of RepSox on prostanoid and thromboxane production in A549 cells. While previous literature extensively described the inhibitory role of PGE_2_ on NK cell cytotoxicity, the role of PGF_2α_ and thromboxane A2/B2 remains poorly understood. High levels of circulating TXB_2_, i.e., the stable metabolite of TXA_2_, are associated with tumor progression, and PGF_2α_ may play a supporting role in tumor progression.^69^ While COX-1 is expressed constitutively, IL-1β was used to induce COX-2 expression in A549 cells, thereby leading to an increase in levels of PGE_2_, PGF_2α_, and TXB_2_ (TXA_2_) (Figures 5F-5H and S4). Dexamethasone suppresses phospholipase A2 (PLA2) expression and activation, therefore reducing AA release from phospholipids and thereby lowering the levels of prostaglandins and thromboxane.^70–72^ This influence is partially reversed by the addition of AA, bypassing the need for PLA2 activity. The COX-2-specific inhibitor Rofecoxib suppresses prostaglandin synthesis, both in the presence and absence of AA, which is in line with its binding mode.^73^ RepSox attenuated the production of PGE_2_, PGF_2α_ as well as TXB_2_/TXA_2_ in the presence of IL-1β (Figure 5E-5H). This effect was rescued by the addition of AA, indicating that RepSox may directly modulate COX-1 or 2 activity in a reversible manner. COX-2 protein expression was detected in A549 cells after stimulation with IL-1β. The addition of Dexamethasone reduced COX-2 levels, whereas, Rofecoxib and RepSox did not change the COX-2 amount (Figure S7). In a biochemical assay, 20 µM RepSox inhibited COX-1 by 61 %, whereas COX-2 was only slightly reduced (13 % inhibition, see also Figure S4). Research by N. Kundu *et al.* demonstrated that inhibiting COX-1 or COX-2 leads to decreased major histocompatibility complex (MHC) class I expression on tumor cells, increasing sensitivity to NK cell-mediated cytolysis.^74^ Simultaneous TGFβR-1 inhibition and alleviation of MHC-I-mediated NK cell inhibition through RepSox may explain our findings.

To characterize the RepSox-induced reversal of the exhausted phenotype of NK cells caused by the artificial tumor microenvironment in an orthogonal setup, we analyzed NK surface receptors by flow cytometry using two antibody panels. These panels covered different surface receptors with various roles for NK cell function or exclusively focusing on NK cell activating receptors, respectively (Figure 6 and S6). The first panel characterized the immunophenotype of NK cells induced by RepSox treatment for the expression of 19 NK cell surface antigens that are related to activation, adhesion, characterization, cytotoxicity, and inhibition of NK cells (Figure 6A). The immunophenotype of unstimulated NK cells is fairly different from the phenotype of NK cells stimulated with IL-15 (Figure 6B). In contrast, NK cells treated with the cytokine mix resemble the unstimulated state. The addition of RepSox to IL-15 did not change NK cell distribution significantly. However, addition of RepSox to the cytokine mix shifted the phenotype in a manner comparable to the IL-15 state. Analysis of the expression pattern of 19 surface antigens in the different conditions led to the emergence of five clusters (Figure 6C). Whereas Cluster 1 represented NK cells with an inactive or suppressed immunophenotype, Cluster 2-5 represented an active NK immunophenotype. Similar to the phenotypic analysis, IL-15 stimulation with or without RepSox decreased the frequency of cluster 1 and increased the frequency of the activated clusters 2-5. The cytokine mix blocked this effect of IL-15, which could be reversed by the addition of RepSox.The surface receptors 41BB and HLA-DR have an important role in the activation of NK cells, while TRAIL is important for cytotoxicity.^75–77^ The expression of 41BB, HLA-DR, and TRAIL was low in the unstimulated condition, however, IL-15-stimulation upregulated their expression (Figure 6D). RepSox addition to IL-15 did not have any effect on their expression. The cytokine mix downregulated of 41BB, HLA-DR, and TRAIL to the level of the unstimulated state. This is in line with studies demonstrating that TGFβ and PGE_2_ can downregulate HLA-DR and TRAIL.^24,78–80^ Addition of RepSox to the combination of the cytokine mix recovered the expression levels. A similar trend was observed for the inhibitory surface receptors TIM3, CD161, and TIGIT, as well as CD11a, which is important for NK cell adhesion (Figures 6D and S6). Analysis of the NK activating receptors NKp30, NKp44, NKp46, 2B4, CD16, DNAM-1, and NKG2D using the same conditions revealed that NKp30, NKG2D, and DNAM-1 protein levels were affected in the same manner (Figure S6H). This confirms previous findings that TGFβ and PGE_2_ can decrease the expression of activating surface receptors such as NKG2D and NKp30.^26^ Additionally, our findings demonstrate that RepSox abrogates the immunosuppressive effects of PGE_2_ and TGFβ on NK cells and restores the expression levels of NK cell surface antigens to the extent of IL-15 treatment. As the addition of RepSox to IL-15 did not have an additional impact on the abundance of NK cell surface antigens, RepSox activity can be narrowed down to the modulation of TGFβ and PGE_2_ signaling.

**Figure 6:**
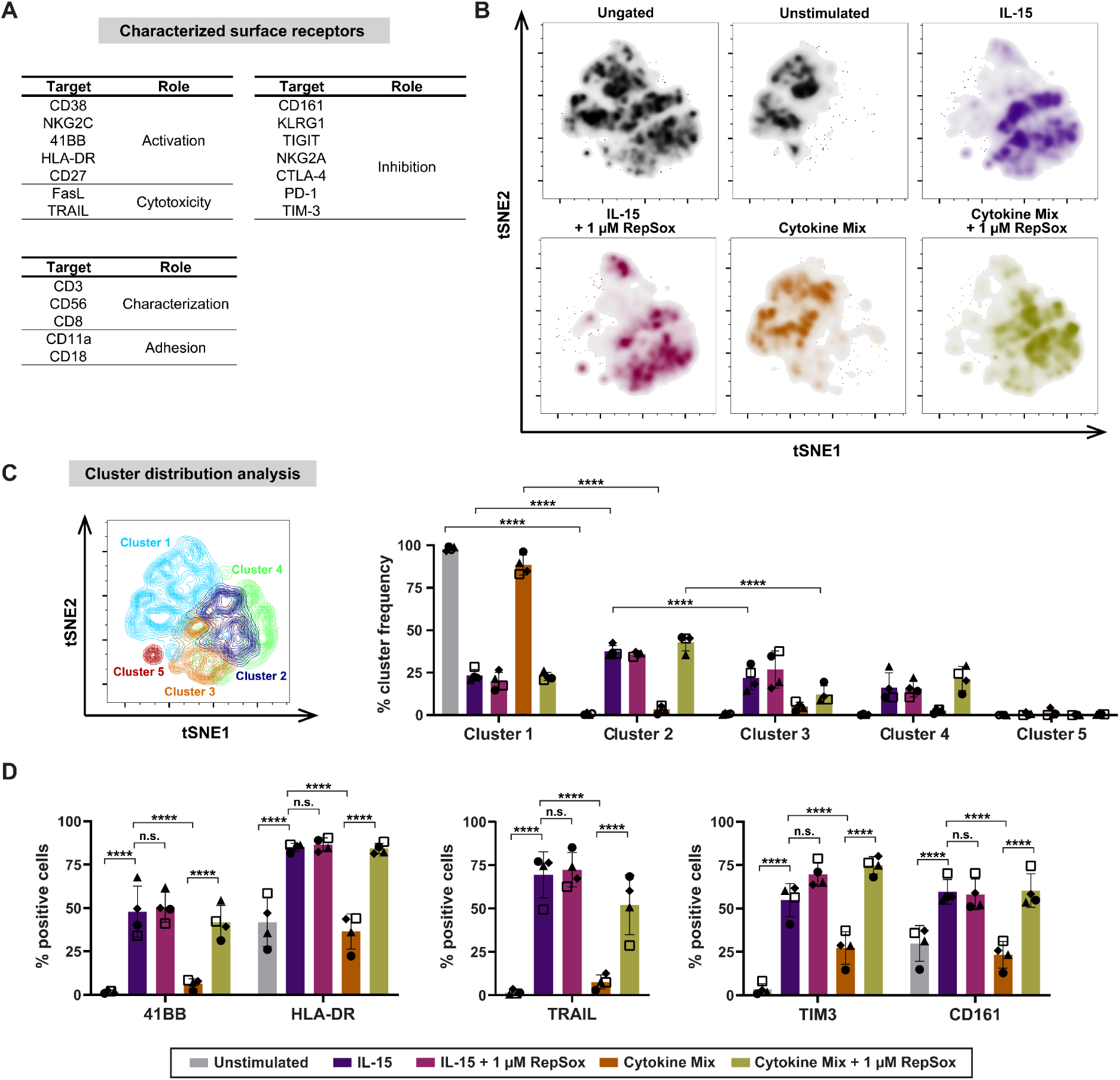
Characterization of NK cell functions upon RepSox-treatment. NK cell surface proteins playing a role in their activation, cytotoxicity, inhibition, adhesion and identity were stained. Lymphocytes of four donors were treated with IL-15 or the cytokine mix individually or in combination with RepSox for 48 h. After staining for various surface proteins, cells were subjected to flow cytometric analysis. Samples were gated on live, single NK cells (CD3^-^ and CD56^+^). The functional markers were analyzed within the whole NK population. The depicted data are mean values of four biological replicates (n=4). A) List of analyzed NK cell functional surface markers. B) The data was submitted to dimensionality reduction using t-distributed stochastic neighbor embedding (t-SNE). C) Cells were clustered based on surface marker expression profiles using the FlowJo™ plugin PhenoGraph algorithm. Cluster frequency is shown for the different conditions. D) Quantified protein expression levels of selected functional markers. Data are mean values ± S.D (n=4); P < 0.05; **, P < 0.01; ***, P < 0.001; ****, P < 0.0001, n.s.: not significant (two-way ANOVA).

## Conclusion

We successfully developed a TME-mimicking phenotypic screening assay for NK cell activity that monitors NK cell-mediated cancer cell killing in a time-resolved manner. The assay enables the co-culture of cancer and NK cells over long incubation times without separate incubation of the cellular components or additives. In a medium-throughput screening, several hit compounds were identified to relieve the suppressive influence on NK cells by different mechanisms of action. Particularly, the TGFβR-1 inhibitor RepSox stood out as it reversed the suppressive influence of both TGFβ and PGE_2_, demonstrating that RepSox has activity beyond TGFβR-1 inhibition that is associated with simultaneous inhibition of COX-1. The here reported assay presents a new approach for the identification of novel small molecule enhancers of NK cell cytotoxicity that contribute to understanding the suppressive mechanisms within the TME in more detail and may lead to the identification of new targets and potential compounds for cancer immunotherapies.

## Author contributions

H W., C.W., and S.Z. designed and guided the research. C.P. adapted the conditions for the NK cell-mediated cell cytolysis assay to a medium-throughput format and carried out the medium-throughput experiments. A.B., E.H., S.K., J.N., A.R., A.K., and D.T. performed the biological experiments. S.S. analyzed the screening data. A.B., E.H., S.Z., and H.W. prepared the manuscript. All authors discussed the results and commented on the manuscript.

## Acknowledgments

This research was supported by the Max Planck Society and was co-funded by the European Union (Drug Discovery Hub Dortmund (DDHD), EFRE-0200481) and Innovative Medicines Initiative (grant agreement number 115489) resources of which are composed of financial contribution from the European Union’s Seventh Framework Programme (FP7/2007-2013) and EFPIA companies’ in-kind contribution. The Compound Management and Screening Center (COMAS) in Dortmund is acknowledged for performing the screening. Dr. Malte Gersch and Nikolas Klink are acknowledged for providing the VHL ligand derivative for testing. A. Binici and E. Hennes would like to acknowledge the International Max-Planck Research School for Living Matter (IMPRS-LM) for their doctoral scholarships. We want to thank Dr. Michael Schulz for his assistance with fluorescence-activated cell sorting. A. Binici would like to express her gratitude to the Joachim-Herz-Foundation for the add-on fellowship.

## Methods

### Cell culture

A549 cells (DSMZ#107) were stably transfected with a linearized pEGFP-B1-H2BJ (Clontech, USA) construct and sorted according to their fluorescence intensity using a BD FACSAria™ Fusion flow cytometer. The resulting A549^Green^ cells were cultured in RPMI-1640 (PAN, #P04-16500) medium supplemented with 10 % heat-inactivated (hi) FBS (Gibco, #10270-106), 1 mM sodium pyruvate (PAN, #P04-43100) and MEM non-essential amino acids (NEAA, PAN, #P08-32100) using 0.7 mg/mL geneticin (G418 disulfate, Sigma Aldrich #G8168). HEK293T cells (ATCC #CRL1268) were cultured in DMEM (PAN, #P04-03550) supplemented with 10 % hi FBS, 1 mM sodium pyruvate, and MEM NEAA. All cultured cell lines were maintained in a humidified atmosphere and 37°C and 5% CO_2_. Cell cultures were regularly analyzed for mycoplasma contamination using the MycoAlert™ Mycoplasma detection Kit following the manufacturer’s protocol (Lonza, #LT07-318) and were always tested negative for mycoplasma infections

### Isolation of human peripheral blood mononuclear (PBMCs) and Natural Killer (NK) cells

Peripheral blood mononuclear cells were isolated from buffy coats of healthy donors (Deutsches Rotes Kreuz (DRK) Hagen, Germany) using Lymphocyte Separating Medium Tubes (PAN, #P04-60125). The buffy coats were diluted in PBS (1:2) and gently layered in the Pancoll-containing tubes. The PBMC layer was separated *via* density centrifugation at 800 x g for 25 min with brake-off. Subsequently, PBMCs were collected, and the remaining red blood cells were removed using ammonium-chloride-potassium lysis buffer (Gibco, #A10492-10). After two additional washing steps using PBS, the purified PBMCs were cryopreserved in hi FBS supplemented with 10% DMSO.

Human NK cells were purified from isolated PBMCs using Dynabeads™ Untouched™ Human NK Cells Kit (Invitrogen™, #11349D) according to the manufacturer’s instructions.

### Manual NK cell-mediated cancer cell cytolysis assay

Cryopreserved donor PBMCs (1*10^6^ cells/mL) were thawed in IMDM medium and placed in tissue culture flasks to isolate the non-adherent lymphocyte fraction. A549^Green^ cells (3,000 cells/well) were then seeded in 96-well plates in phenol red-free RPMI1640 medium, and the cells were incubated overnight at 37°C with 5% CO_2_. The next day, lymphocytes or NK cells were added together with IL-15 (30 ng/mL for lymphocytes and 7.5 ng/mL for NK cells), TGFβ-1 (30 ng/mL for lymphocytes and 7.5 ng/mL for NK cells) and PGE_2_ (200 ng/mL) in accordance to the previously assessed effector-to-target (E:T) ratio.

Subsequently, compounds or DMSO were added to the samples that were treated with cytokine mix (IL-15, TGFβ-1 and PGE_2_) samples and A549^Green^ cell count was monitored using kinetic live-cell analysis in 4 h intervals using the IncuCyte S3 device at 10X magnification. Data analysis was performed using the IncuCyte® S3 2019B Rev software. To calculate the cytolysis rate, the area under the curve (AUC) was determined using GraphPad Prism 9, and values were normalized to the values for cells that treated with IL-15 (set to 100% cytolysis) and cytokine mix (set to 0% cytolysis). The half-maximal effective concentration (EC_50)_ values were determined using non-linear regression *via* a four-parameter fit using GraphPad Prism 9 (GraphPad Software, USA).

### Automated NK cell-mediated cancer cell cytolysis assay

Previous to the screening, lymphocytes from different donors were assayed to determine the ideal E:T ratio for achieving the highest S/B value. For the high-throughput screening, A549^Green^ cells (500/well for 384-well plate and 400 cells/well for 1536-well plates) were seeded in phenol red-free RPMI1640 using the Multidrop Combi Device (Thermo Fisher Scientific) and incubated overnight at 37°C and 5% CO_2_. Compound treatment was conducted using an Echo550 dispenser (Labcyte) using a final concentration of 11 µM or a serial dilution to determine EC_50_ values. Subsequently, the lymphocyte suspension was prepared, combined with IL-15 (30 ng/mL) and the inhibitory factors TGFβ-1 (30 ng/mL) and PGE_2_ (200 ng/mL) and then added to the screening plate using the Multidrop Combi (Thermo Fisher Scientific).

Untreated and IL-15-treated lymphocytes were used as control samples. Imaging of the A549^Green^ cells was carried out using the ImageXpress Micro XL device (Molecular Devices) at 10x magnification (excitation: 465/40 nm, emission: 525/30 nm). Image analysis was carried out using the Cell Proliferation HT module of MetaXpress. The A549^Green^ count was determined in relation to the cytokine mix and DMSO-treated co-culture.

### End-point evaluation of IFN-γ secretion levels

Subsequent to the NK cell-mediated cancer cell cytolysis assay, supernatants were harvested and assayed for IFN-γ levels using the ELISA MAX™ Standard Set Human IFN-γ (Biolegend, #430101) according to the manufacturer’s instructions.

### Cell growth and viability

A549^Green^ cells (3,000 cells/well in 96-well plates) were seeded and allowed to attach overnight at 37°C and 5% CO_2_. The next day, cells were treated with cytokines, compounds or DMSO as a control and subjected live-cell analysis for 96 h using the IncuCyte ZOOM or S3 live-cell imaging system (Sartorius). The cell count was obtained from the time-lapse image acquisition using the IncuCyte ZOOM (2018A) or IncuCyte S3 (2022B) software.

### *SBE4* promoter-dependent reporter gene assay

HEK293T cells (40,000 cells/well in 96 well plates) were reverse-transfected with a *SBE4*-dependent firefly luciferase reporter (SBE4-Luc, Addgene, #16495) and a constitutively expressed *Renilla* luciferase reporter(pRL-TK-Rluc, Promega) using Lipofectamine™ 2000 (Invitrogen™, #11668019) and RPMI-1640 medium with reduced serum concentration (2 % hi FBS).^59^ Cells were allowed to attach for 6 h and subsequently treated with 20 ng/mL TGFβ-1 and compounds or DMSO for 24 h. A control without TGFβ-1 stimulation was included. The firefly (Fluc) and *Renilla* (Rluc) luciferase luminescence were determined by means of the Dual-Glo® Luciferase Assay System (Promega, #E2940) using the TECAN Spark® Multimode Microplate Reader. The *SBE4* promoter activity was determined by normalizing the Fluc signals to the corresponding Rluc signals and setting the ratio of the TGFβ-1/DMSO-treated cells to 100 %. Dose-response curves and the half maximal inhibitory concentration (IC_50_) values were obtained and fitted using GraphPad Prism 9.0 (GraphPad Software, Inc, USA) using a variable slope non-linear regression four-parameter curve fit.

### Prostaglandin formation assay

A549 cells (1.5*10^6^ cells/dish) were seeded using DMEM medium and incubated overnight. The next day, cells were treated with serum-reduced DMEM containing 2 % FBS (v/v) with or without 1 ng/mL IL-1β and compounds or DMSO and incubated for 16 h. This was followed by collecting the supernatants and the assessment of the prostanoid and thromboxane levels by LC-MS/MS.^81^

### Enzymatic cyclooxygenase 1 and 2 activity assay

The enzymatic assays to assess the activity of human recombinant cyclooxygenase 1 and 2 were conducted by Eurofins Discovery. For COX-1, 3 µM arachidonic acid, 25 µM 10-acetyl-3,7-dihydroxyphenoxazine (ADHP), and 20 µM RepSox were incubated for 3 min at room temperature. For COX-2, 1.2 µM arachidonic acid, 25 µM ADHP, and 20 µM RepSox were incubated for 5 min at room temperature. Enzymatic activity was measured using fluorimetric detection with resorufin oxidized by HRP.

### Immunoblotting

Lymphocytes (2,000,000 cells/mL) were stimulated with 30 ng/mL IL-15, 30 ng/mL TGFβ-1 and 200 µg/mL PGE_2_, and treated with either compound or DMSO for 48 h. For granzyme B detection, total cellular protein was extracted using ice-cold NP-40 alternative-based buffer (50 mM sodium chloride, 50 mM TRIS, and 1 % NP-40 alternative, pH 8.0) supplemented with PhosSTOP™ phosphatase (Roche) and cOmplete™ protease inhibitor cocktail (Roche)). Briefly, after the removal of cell debris *via* centrifugation at 20,000xg for 25 min, the protein concentration was determined by means of the DC protein assay (Bio-Rad Laboratories), followed by the addition of 1X SDS loading buffer to the samples. Proteins were separated using 10 % polyacrylamide gel and transferred to a polyvinylidene difluoride (PVDF) membrane using wet tank transfer in blotting buffer (192 mM glycine, 25 mM TRIS, 10 % methanol). Next, membranes were blocked using LI-COR Intercept® PBS blocking buffer for 60 min at room temperature. Afterwards, the membranes were incubated with the primary antibodies overnight at 4°C. To detect granzyme B, the respective primary antibodies (Granzyme B 1:2,000, ab134933, Abcam, UK) and vinculin (1:5,000, V9131, Merck KGaA, DE) were mixed with LI-COR Intercept® PBS blocking buffer. The proteins were visualized using secondary antibodies coupled to IRDye® 800CW (LI-COR Biosciences, US) and bands were detected utilizing the ChemiDoc MP Imaging System (Bio-Rad Laboratories, DE). Band intensities were quantified using the ImageLab software (Bio-Rad). Granzyme B protein levels were normalized to the levels of the reference protein vinculin, and the IL-15 condition was set to 100%.

For detection of COX-2, A549 cells were harvested after the prostaglandin formation assay and lysed in hot (96°C) SDS buffer (56 mM Tris-HCl pH6.8 buffer with 2.2 % SDS and 11 % glycerol) supplemented with protease inhibitors (cOmpleteTM Mini, Roche). Afterwards, the lysates were sonicated (low pulse, 3x 10 s) to remove the DNA jelly formed during lysis. Then, total protein concentration was measured using the PierceTM BCA Protein Assay Kit on an infinite M200 microplate reader (Tecan Group Ltd., Männedorf, Switzerland). Next, cell lysates were separated by SDS-polyacrylamide gel electrophoresis followed by Western Blot transfer to nitrocellulose membranes (0.2 µm, Bio-Rad, Hercules, CA, USA) using the Trans-Blot Turbo Transfer System (Bio-Rad, Hercules CA, USA). The Precision Plus Protein All Blue Prestained Protein Ladder (Bio-Rad, Hercules CA, USA) was used as a molecular weight marker. After blotting, the membranes were blocked with EveryBlot blocking Buffer (Bio-Rad, Hercules CA, USA; 1 h, RT). Then, membranes were incubated with primary antibodies directed against COX-2 (sc-19999, Santa Cruz Biotechnology, USA) and α-tubulin (housekeeping control, sc-5286, Santa Cruz Biotechnology, USA) followed by incubation with the respective fluorescence-conjugated secondary antibodies (IRDye, LI-COR Biosciences, Bad Homburg, Germany). Next, the membranes were scanned with the Odyssey Infrared Imaging System (LI-COR Biosciences, Bad Homburg, Germany) and immunoreactive bands were quantified employing the Image Studio 5.2 software (LI-COR Biosciences, Bad Homburg, Germany).

### Flow cytometry-based characterization of lymphocytes

The T cell and NK cell-specific surface markers CD3 and CD56 were stained in order to determine the proportion of NK cells in donor lymphocytes or purified NK cells. For this purpose, lymphocytes or purified NK cells were resuspended in FACS buffer (PBS supplemented with 2 % FBS) and anti-CD3-PE (1:200, BioLegend, #300308) and anti-CD56-BV421 (1:100, BD Bioscience, #562751) antibodies were added, followed by a 20-min incubation on ice. Afterwards, the cells were washed twice using FACS buffer and resuspended in FACS buffer. Next, the dead-cell stain 7-AAD (1:50) was added, and the samples were measured using a BD-LSR II flow cytometer. The samples were excited by utilizing a 405 nm and a 488 nm laser and emission was detected by utilizing 450/50, 575/26 as well as 695/40 nm filters. Data analysis was performed by using FlowJo™ 10.7.2.

To characterize NK cells, stimulated lymphocytes were stained with two different panels: a mixed receptor panel and an activating receptor panel. Cell counts were determined using a CASY Cell Counter & Analyzer (OLS® OMNI Life Science, Bremen) and adjusted. First, cells were stained with Zombie NIR (BioLegend (San Diego)) diluted 1:700 in PBS and afterwards washed with FACS-buffer. Antibody dilutions were prepared in FACS buffer in the presence of 50 % Brilliant Stain Buffer (BD Bioscience (San Jose)) and cells stained. The mixed receptor panel included the following antibodies: CD38(HB7)-BUV395 (BD Bioscience (San Jose)), NKG2C(134591)-BUV496 (BD Bioscience (San Jose)), CD3(UCHT1)-BUV563 (BD Bioscience (San Jose)), CD161(HP-3G10)-BUV661 (BD Bioscience (San Jose)), CD56(B159)-BUV805 (BD Bioscience (San Jose)), 41BB-(4B4-1)-BV421 (BioLegend (San Diego)), KLRG1(2F1)-BV510 (BioLegend (San Diego)), HLA-DR(L243)-BV605 (BioLegend (San Diego)), FasL(NOK-1)-BV650 (BD Bioscience (San Jose)), TIGIT(741182)-BV786 (BD Bioscience (San Jose)), CD11a(HI111)-AF488 (BioLegend (San Diego)), CD8a(RPA-T8)-AF532 (Thermo Fischer Scientific (Waltham)), CD27(M-T271)-PerCP-Cy5.5 (BD Bioscience (San Jose)), NKG2A(Z199)-PE (Beckman Coulter (Brea)), CTLA-4(BNI3)-PE-Dazzle (BioLegend (San Diego)), TRAIL(N2B2)-PE-Cy7 (BioLegend (San Diego)), PD-1(EH12.1)-AF647 (BD Bioscience (San Jose)), CD18(TS1/18)-AF700 (BioLegend (San Diego)) and TIM-3(F38-2E2)-APC-Fire 750 (BioLegend (San Diego)). While the activating receptor panel contained following antibodies: CD3(UCHT1)-BUV563 (BD Bioscience (San Jose)), CD56(B159)-BUV805 (BD Bioscience (San Jose)), NKp46(9E2)-BV421 (BioLegend (San Diego)), 2B4(C1.7)-FITC (BioLegend (San Diego)), NKp44(P44-8)-PerCP-Cy5.5 (BioLegend (San Diego)), CD16(3G8)-PE-Dazzle (BioLegend (San Diego)), DNAM-1(DX11)-AF647 (BD Bioscience (San Jose)), NKG2D(FAB139N)-AF700 (R&D Systems (Minneapolis)) and NKp30(P30-15)-APC-Fire750 (BioLegend (San Diego)). Cells were then washed twice with FACS buffer and analyzed using a Cytek® Aurora (5L, Cytek Bioscience Inc., Fremont) with the SpectroFlo® 3.1 software. Data was analyzed with FlowJo™ 10.8.2. The FlowAI 2.3.1 plug-in was used at default settings to clean the data. Afterwards, manual gating was performed to reach NK cells. Percent positive cells or geometric mean fluorescence intensity (gMFI) was analyzed to best fit the populations. Additionally, to perform dimensionality reduction, an equal number of NK cells per donor and stimulation condition were concatenated, and the FlowJo™ inbuild tSNE algorithm was used at default settings. Cluster analysis was carried out using the PhenoGraph clustering algorithm.

### Statistical analysis

If not stated otherwise, experiments were performed with at least three biological replicates (n≥3) and two technical replicates (N≥2) per biological replicate. Experiments performed with primary lymphocytes or purified NK cells were conducted with at least three donors, where donors vary between experiments. Various symbols were utilized to depict the activity of each donor. The error bands in the graphs indicate the standard deviation of the data set. One- or two-way ANOVA were used to determine the significance of different treatments and performed using GraphPad Prism 9 (GraphPad Software, Inc, USA).

