## Supplementary Information for "Identification of small molecule enhancers of NK cell tumoricidal activity via a tumor microenvironment-mimicking co-culture assay"

### Supporting Figures

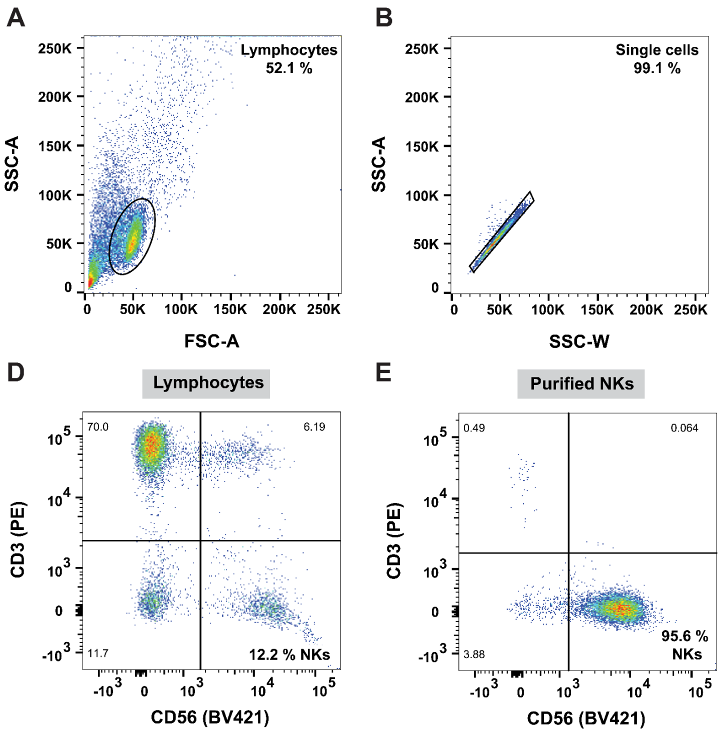


**Figure S1: FACS-based determination of NK cell proportions.** Lymphocytes or purified NK cells were stained for CD3, CD56, and 7-AAD for live/dead discrimination and subjected to flow cytometry-based analysis. A)-C) Utilized gating strategy. A) Forward scatter was plotted against side scatter to identify the lymphocytes subset. Next, doublets were excluded using SSC-A vs. SSC-W (B), and finally, only live cells were further analyzed for CD3 and CD56 expression (C). D) and E) Analysis of CD3 and CD56 expression levels for donor lymphocytes and NK cells, respectively. The NK cell proportion, i.e., CD3-/CD56+, is represented in the bottom right quadrant.

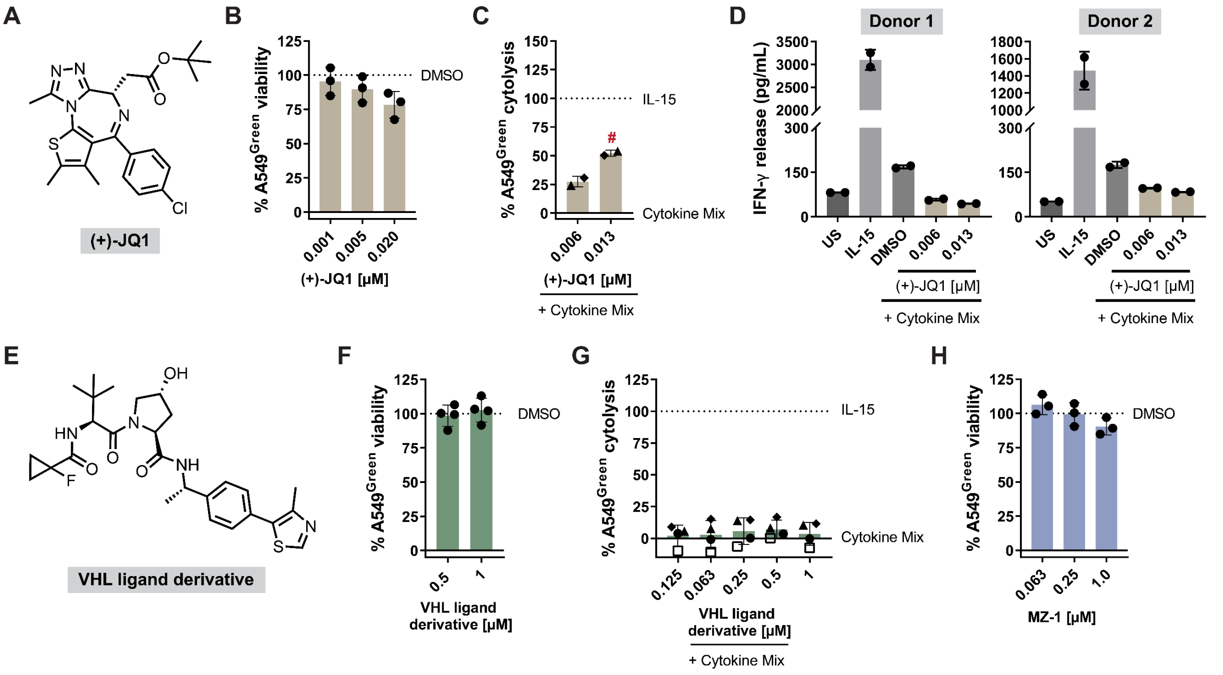


**Figure S2: Influence of (+)-JQ1 and a VHL ligand derivative on NK cell cytotoxicity.** A) Chemical structure of (+)-JQ1. B) A549$^{Green}$ cells were treated with (+)-JQ1 for 96 h. The viability was determined using the AUC of obtained cell count data and normalized to their corresponding DMSO control, which was set to 100%. Data are mean values ± SD (n=3). C) NK cell-mediated cancer cell cytolysis assay using lymphocytes in the presence of (+)-JQ1. Data are mean values ± SD (n=2, N=3 for (+)-JQ1). # = condition reduces cell viability. D) IFN-γ levels. Supernatants of NK cell-mediated cancer cell cytolysis assays using lymphocytes were collected after 144 h, followed by assessing IFN-γ levels. Data are mean values ± SD (N=2, n=3). E) Chemical structure of the VHL ligand derivative. Data are mean values ± SD (n=4). F) Viability of A549$^{Green}$ cells treated with VHL ligand derivative for 96 h. G) NK cell-mediated cancer cell cytolysis assay using lymphocytes in the presence of the VHL ligand derivative. Data are mean values ± SD (N=3, n=4 for the VHL ligand derivative). H) Viability of A549$^{Green}$ cells treated with MZ-1 for 96 h.

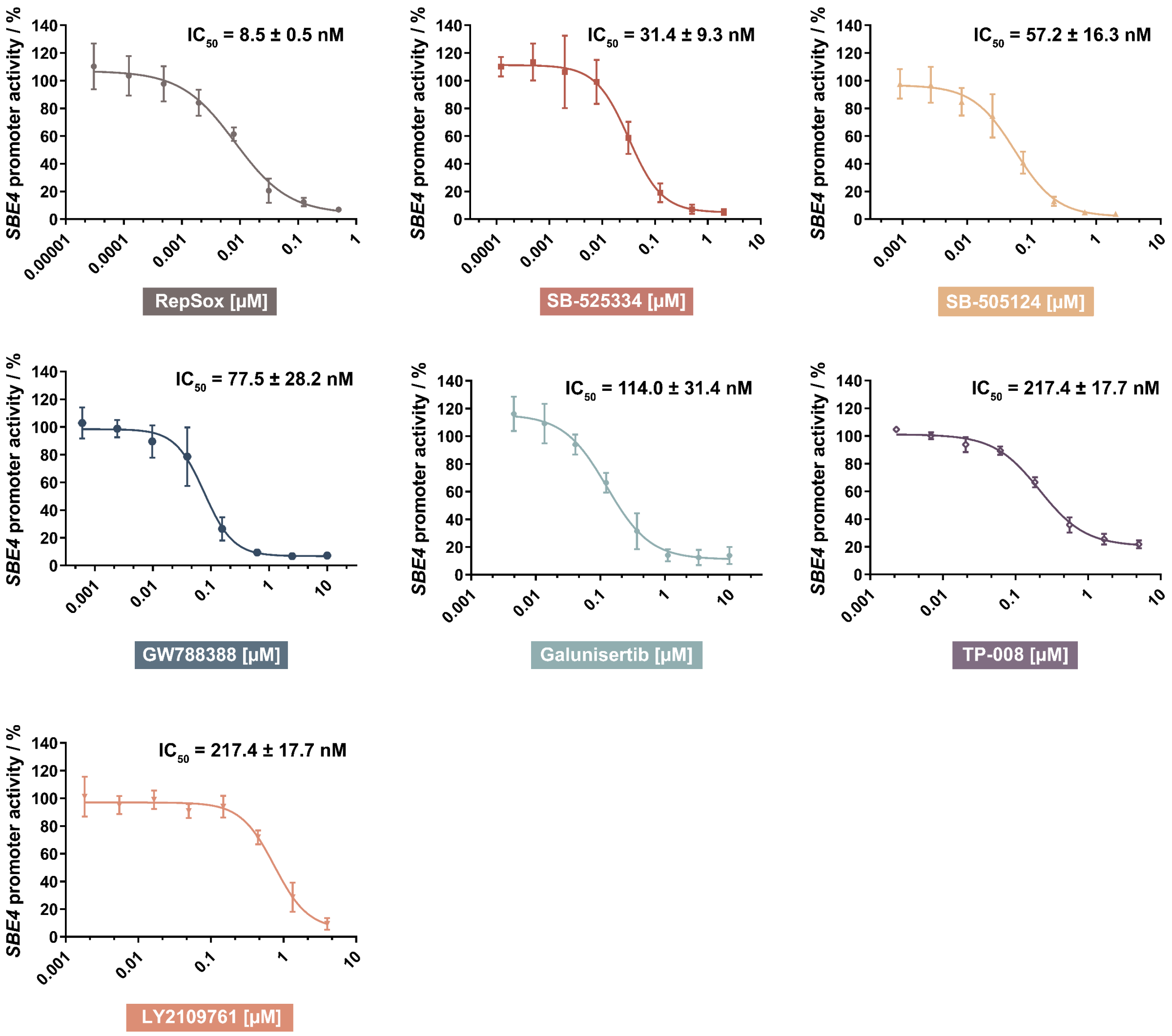


**Figure S3: Comparison of the inhibitory effect of TGFβR-1 inhibitors on the *SBE4*-dependent reporter activity.** HEK293T cells were transiently transfected with a *SBE4* promoter-controlled firefly luciferase (Fluc) and a *Renilla* luciferase (Rluc) construct followed by TGFβ stimulation to upregulate Fluc expression and simultaneous compound treatment for 24 h. Fluc data were normalized to the corresponding Rluc data. Data are mean values ± SD (n=4 for RepSox, SB-525334 as well as galunisertib, n=3 for SB-505124, GW788388 and LY2109761). Related to Figure 4B.

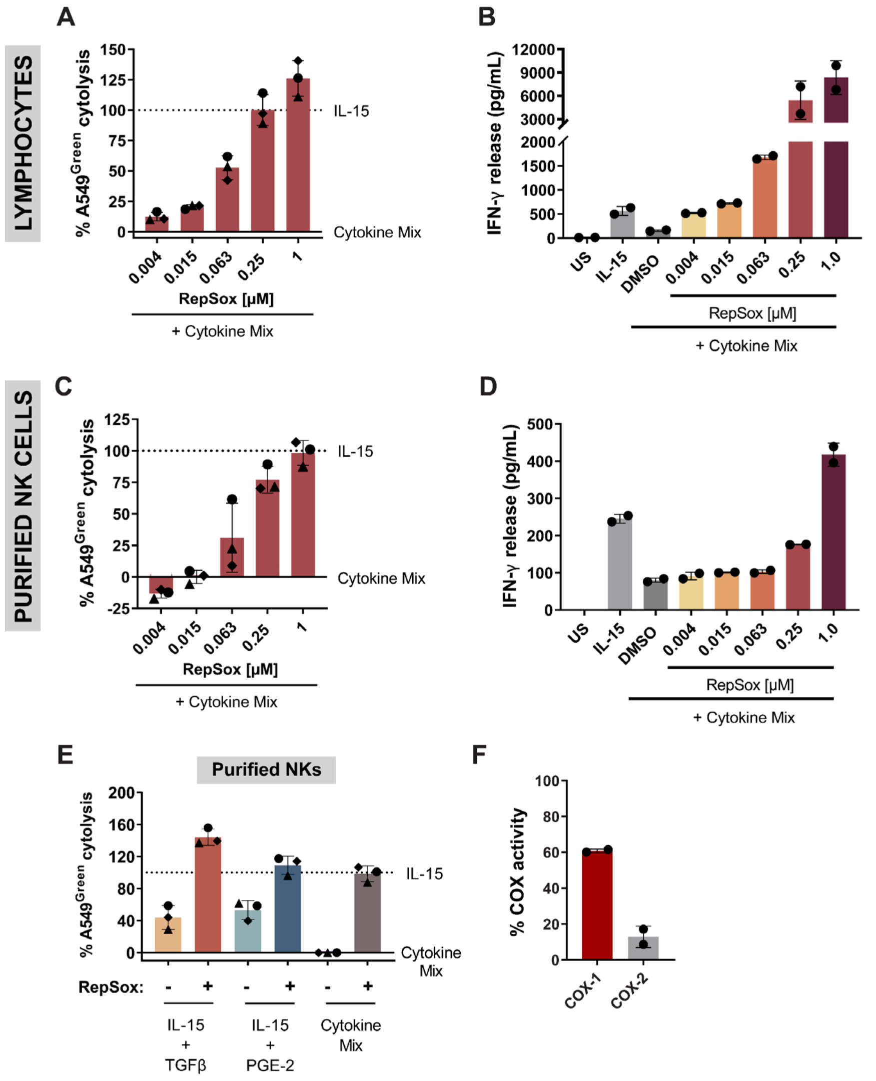


G

| Condition | $PGE_2$ [pg/mL] | $PGF_{2\alpha}$ [pg/mL] | $TXB_2/TXA_2$ [pg/mL] |
|---|---|---|---|
| DMSO | 735.8 ± 203.6 | 264.9 ± 78.7 | 45.7 ± 10.5 |
| RepSox 1 µM | 505.7 ± 95.9 | 173.9 ± 76.5 | 11.8 ± 20.4 |
| RepSox 5 µM | 389.4 ± 106.3 | 128.2 ± 57.3 | 17.9 ± 15.5 |
| Dexamethasone | 13.3 ± 18.8 | - | - |
| Rofecoxib | - | - | - |

**Figure S4: RepSox enhances NK cell cytotoxicity.** A) NK cell-mediated cancer cell cytolysis assay using lymphocytes treated with RepSox. Data are mean values ± SD (n=3). B) Assessment IFN-γ levels in supernatants of NK cell-mediated cancer cell cytolysis assays using lymphocytes treated with RepSox for 144 h. Data are mean values ± SD (N=2, n=3). C) NK cell-mediated cancer cell cytolysis assay using RepSox-treated purified NK cells. Cells were treated for 144 h. Data are mean values ± SD (n=3). D) Assessment of IFN-γ levels in supernatants of NK cell-mediated cancer cell cytolysis assays using RepSox-treated purified NK cells after 144 h. Data are mean values ± SD (N=2, n=3). E) Influence of 1 µM RepSox on the NK cell cytolytic activity using purified NK cells. Data are mean values ± SD (n=3). Related to Figures 4 and 5. F) Influence of RepSox (20 µM) on COX-1/2 activity. Data are mean values ± SD (N=2). G) Mean of the prostanoid and thromboxane release into A549 supernatants. Data are mean values ± SD (N=3).

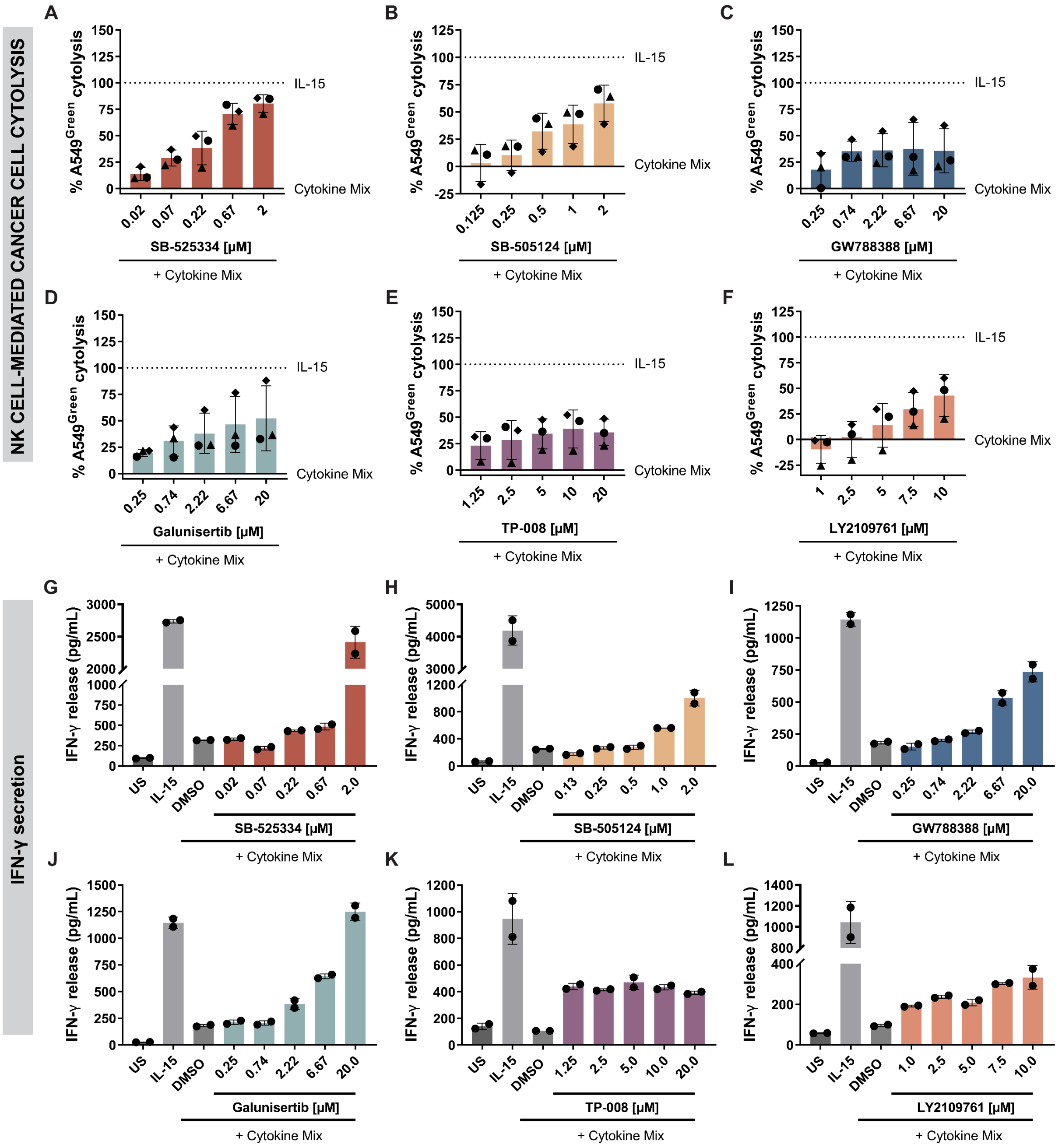


**Figure S5: Comparison of the cytolysis-enhancing activity of TGFβR-1 inhibitors in NK cell-mediated cancer cell cytolysis assay.** A-F) A549$^{Green}$ cytolysis using lymphocytes treated with SB-525334, SB-505124, GW788388, galunisertib, TP-008 and LY2109761. Data are mean values ± SD (n=3, N=3). G-L) IFN-γ levels in supernatants of NK cell-mediated cancer cell cytolysis assays using lymphocytes treated with SB-525334, SB-505124, GW788388, galunisertib, TP-008 and LY2109761 after 144 h. Data are mean values ± S.D. (N=2, n=3). Related to Figure 4.

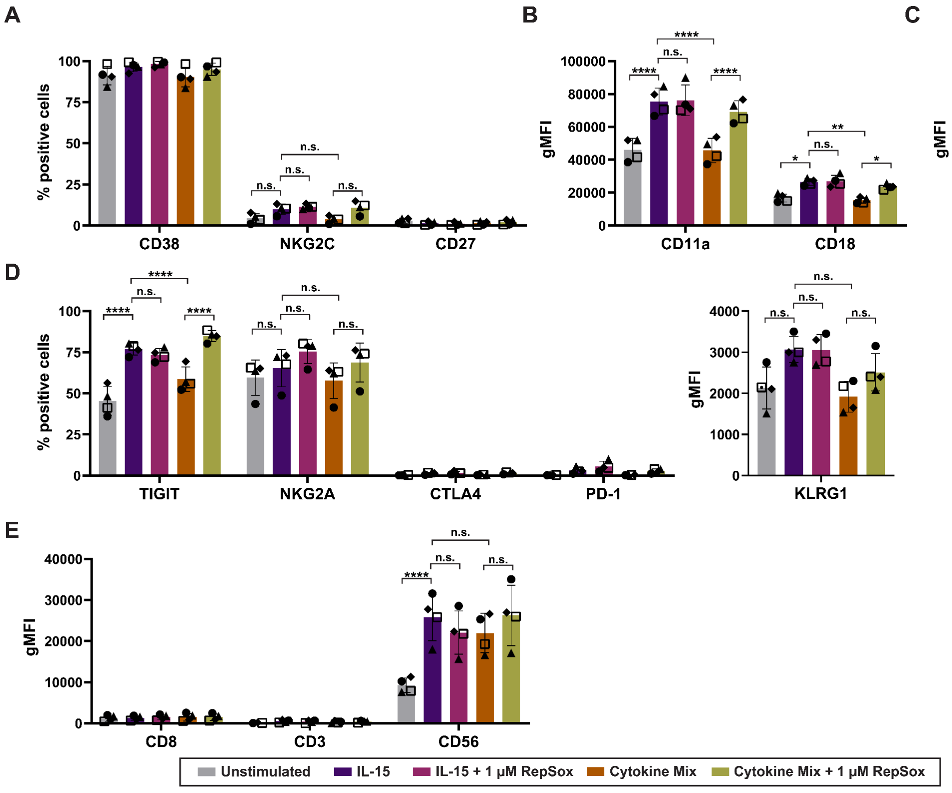


F

**Characterized surface receptors**

| Target | Role |
|---|---|
| NKp30<br>NKp44<br>NKp46<br>2B4<br>CD16<br>DNAM-1<br>NKG2D | Activation |

| Target | Role |
|---|---|
| CD3<br>CD56 | Characterization |

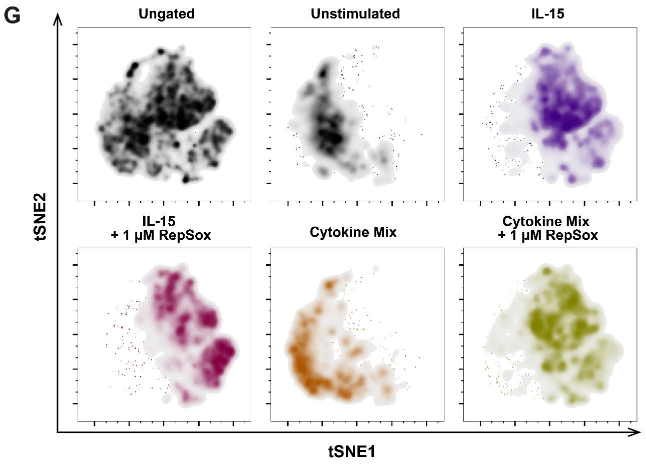


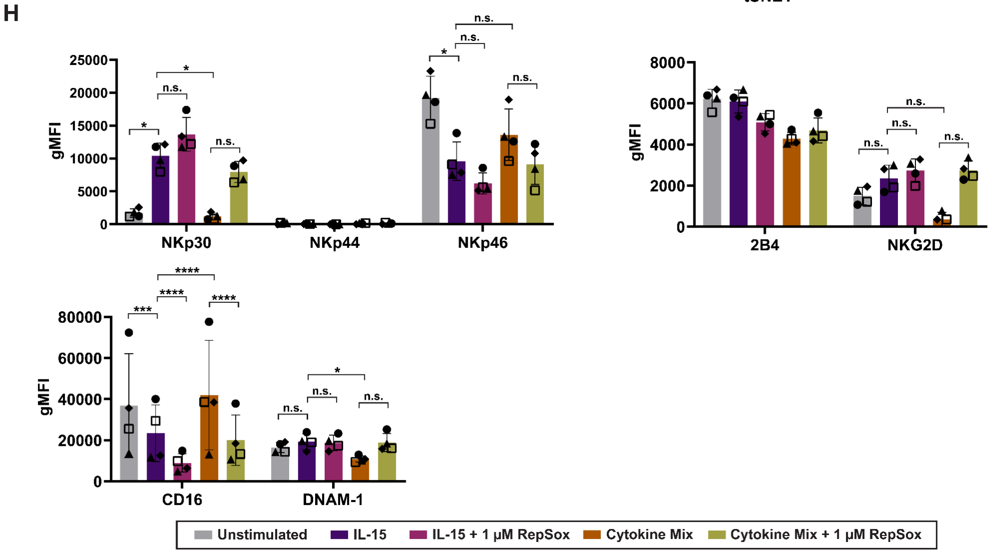


**Figure S6: Characterization of NK cell surface receptor expression upon RepSox treatment.** Lymphocytes were treated with activating IL-15 or the cytokine mix individually or combined with 1 µM

RepSox for 48 h. Subsequently, various NK cell surface marker proteins were stained and quantified using flow cytometry. A-E) Quantified protein levels of selected functional marker proteins. Data are mean values ± S.D (n=4); P < 0.05; **, P < 0.01; ***, P < 0.001; ****, P < 0.0001 (two-way ANOVA). F) List of analyzed activating receptors. G) Data was submitted to dimensionality reduction using t-SNE and then to cluster analysis based on surface marker expression profiles. H) Quantified protein levels of selected functional marker proteins. Data are mean values ± S.D (donors=4); P < 0.05; **, P < 0.01; ***, P < 0.001; ****, P < 0.0001 (two-way ANOVA). Related to Figure 6.

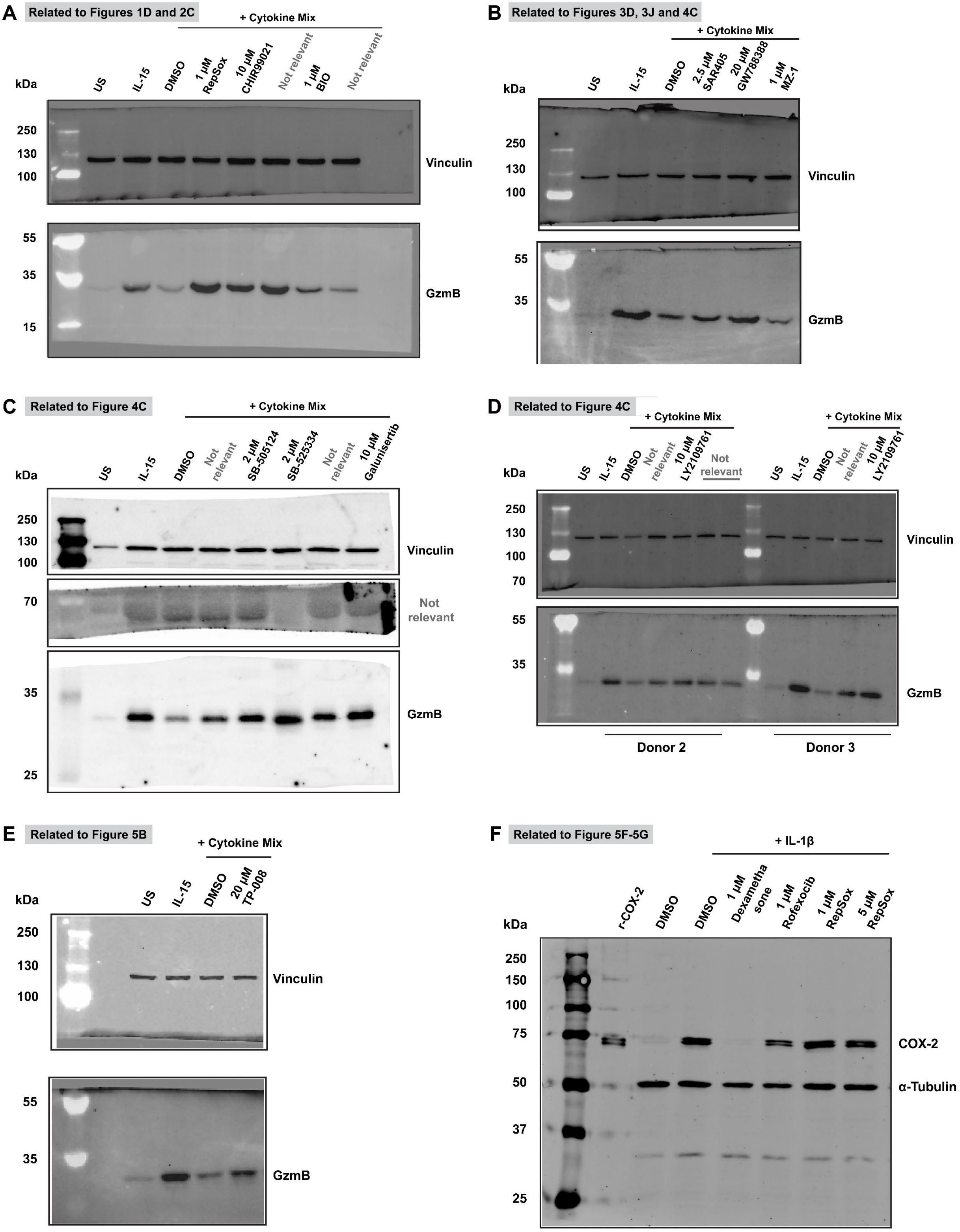


**Figure S7:** A-E) Influence of IL-15, cytokine mix individually or combined with compounds on granzyme B expression. Lymphocytes of three donors were treated using the indicated conditions for 48 h. Uncropped immunoblots are shown. All depicted immunoblots are representative for three biological replicates (n=3), except for SB-505124, for which the immunoblot is representative for n=4. Related to Figures 1-4. F) Influence of RepSox on COX-2 protein levels. A549 cells were treated with IL-1β to induce COX-2 expression and simultaneously treated with RepSox for 16 h. COX-2 expression inhibitor Dexamethasone and COX-2 inhibitor Rofecoxib were used as controls.
